# A Neyman-Pearson Framework for Modeling Cellular Decision Making Using Single-Cell TNF–NF-κB Signaling Data

**DOI:** 10.64898/2025.12.30.696266

**Authors:** Ali Emadi, Tomasz Lipniacki, Andre Levchenko, Ali Abdi

## Abstract

Cells make hard calls under noise. When signaling is abnormal, those calls can go wrong and drive pathological conditions and diseases. In this research, we develop a Neyman–Pearson (NP) detection-theory framework that maximizes probability of detection (*P*_D_) for a chosen false alarm probability (*P*_FA_), without requiring prior probabilities, using experimental single-cell measurements of NF-κB responses to tumor necrosis factor (TNF), a critical pathway involved in cell survival, apoptosis, immune signaling, and stress response, in wild-type and A20-deficient fibroblasts. We model log-responses as (multi)variate Gaussian and compute optimal thresholds, *P*_D_–*P*_FA_ trade-offs, and ROC curves at 30 minutes and 4 hours. The NP framework captures expected biology: *P*_D_ increases with TNF dose; wild-type cells outperform A20^-/-^ at matched conditions; and combining two time points (bivariate analysis) improves detection (e.g., for 0.0052 vs. 0.2 ng/mL, *P*_D_ rises from 0.71 (30 minutes) and 0.42 (4 hours) to 0.80 at *P*_FA_ = 0.1). The analysis recovers expected biology (higher TNF causes higher detectability; negative feedback lowers late responses) and flags cases where decision quality degrades (e.g., perturbations that blunt separation between conditions). The same recipe extends to multivariate readouts without changing the logic.

Overall, the NP detection framework provides a compact, quantitative score of pathway performance and failure. It turns noisy single-cell readouts into actionable decision metrics that compare doses, time points, and perturbations, and ultimately, can help explain when and how cellular decisions drift toward pathology.

## INTRODUCTION

In the field of cellular biology, the signal recognition and response of cells to environmental stimuli are key concepts. These processes are commonly involved in intricate biochemical signaling networks, where the strength and concentration of extracellular signals can influence cell fate and intercellular phenomena [1]-[3]. By comprehending the intricacies of such interactions, we can improve our understanding of cellular behavior, which may lead to novel approaches for the diagnosis and treatment of various pathological conditions. Therefore, it is crucial to study the mechanisms involved in this process to gain deeper insights into cellular decision-making. Molecular noise is a common phenomenon in biological systems [4], and its presence can impede the ability of cells to accurately sense and respond to various input signals [5]. The inherent stochasticity of biological noise renders cellular decision-making probabilistic in nature [5]. A wide range of potential outcomes can impact the cell’s behavior and the system’s overall function, each with varying degrees of likelihood. Even though biological noise presents significant challenges, researchers are actively seeking new ways to reduce its effects and improve the accuracy of cellular signaling and decision-making processes. Specifically, it has been shown that signal transduction noise arises from both extrinsic and intrinsic sources. Intrinsic noise is caused by the stochastic nature of biochemical reactions, which involve a small number of molecules. On the other hand, extrinsic noise results from variations in the levels of TNF receptors and cellular heterogeneity upon stimulation [6]. To gain a better understanding of the probabilistic nature of cellular decision-making, we employ a system model with explicit statistical parameters for alternative outcomes and quantify detection performance from single-cell data. Recently, similar models have used decision theory to derive optimal thresholds and error probabilities from such measurements. The objective of such a model is to assess the system’s capability to accurately detect the presence or absence of the desired signal. In those studies, the prior probabilities for signal presence/absence were assumed to be known, typically equal, that is 0.5/0.5, and then the maximum likelihood decision approach was applied [7]-[9]. It is important to consider a different modeling approach when the prior probabilities of signal presence are unknown. This paper introduces a signal detection model where the cell maximizes the probability of detecting the signal for any given false alarm probability. The signal detection model used in this study is based on the Neyman-Pearson signal detection theorem, which takes into account the probability of falsely detecting a signal when it is absent, also known as the false alarm probability. These quantitative and probabilistic models also have the potential to shed light on the stochastic signaling mechanisms and phenotype induction associated with genetic mutations that cause diseases [10]. By providing a deeper understanding of these mechanisms, our proposed model may contribute to the development of novel approaches for the diagnosis and treatment of genetic disorders. In order to validate such a method in a biologically meaningful context, we utilized a signaling pathway involving the regulation of the transcription factor Nuclear Factor κB (NF-κB) by tumor necrosis factor (TNF) (Fig. 1) [5]. This allowed us to investigate the method’s effectiveness in a real-world scenario and assess its potential for future applications. TNF is a crucial antiviral cytokine that can cause significant damage to healthy host tissues. In addition to its role in activating NF-κB, a critical gene regulator involved in cell survival, viral replication, and pathological processes such as autoimmune diseases, various types of cancer, and inflammation, TNF has also been shown to mediate both anti-apoptotic and pro-apoptotic signals and can even trigger necroptosis, a type of pro-inflammatory cell death [11]. Upon activation, NF-κB can act to inhibit apoptosis in cells. The presence of the A20 molecule can also help reduce noise in the signaling pathway by acting as negative feedback to the information bottleneck. However, the effect of A20 activation on the amount of information attained in the system is variable and can either increase or decrease information [5]. The loss or mutation of the A20 molecule, inhibitory feedback of the NF-κB pathway, has been implicated in various pathological conditions such as cancer and chronic inflammation [12][13]. The negative feedback mediated by A20 can result in the reduction of NF-κB levels, leading to downstream effects on cellular signaling. Also, A20 deficiency may contribute to cancer development and lead to significant pathological conditions. This effect is believed to occur due to the persistent activation of anti-apoptosis genes mediated by NF-κB, as described in [14]. In response to NF-κB activation, the transcription of early genes, including cytokines, can be upregulated [7]. Furthermore, tumor microenvironments often contain heterogeneous cell populations with varying degrees of A20 expression, which makes mixed populations particularly relevant for understanding signal discrimination in cancer contexts.

**Figure 1.**
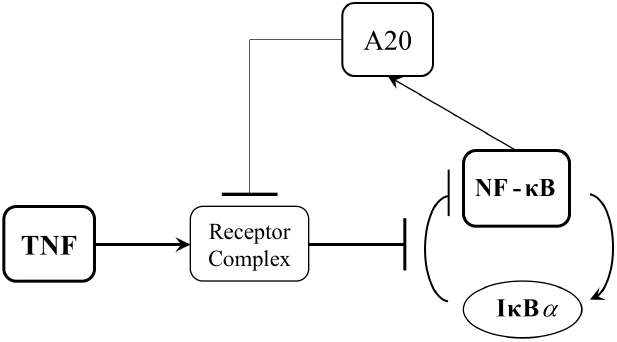
The Role of A20 in TNF– NF-κBsignaling pathway.

The objective of this paper is to demonstrate the applicability of the proposed statistical decision-making framework to diverse molecular signaling systems. We aim to showcase the versatility and effectiveness of this framework across different biological contexts when there is no knowledge of priors. The aim is twofold: first, to observe how the suggested decision model is expanded to examine various time points in the process of signaling outcomes using statistical multi-dimensional analysis for the TNF–NF-κB. Second, using the widely recognized graphical tool, the receiver operating characteristic (ROC), to visualize cell decision-making performance in both normal and abnormal conditions. The overall objective is to determine the probabilities of cell decision-making and establish a correlation between statistical parameters and metrics with biological discoveries.

The rest of the paper is organized as follows. First, introducing the single-cell experimental data of the important TNF–NF-κB signaling pathway, then the dynamic range of NF-κB responses in the form of means and variances of different TNF levels is shown. After that, the decision probabilities of univariate and bivariate decision analysis are presented in RESULTS. Finally, some concluding remarks are listed at the end.

### SINGLE-CELL DATA OF THE TNF–NF-κB SYSTEM

The data set is composed of wild-type and A20^−/−^ (A20-deficient) 3T3-immortalized mouse embryonic fibroblasts, which can also be labeled as normal and abnormal cells, respectively. Nuclear NF-κB concentrations were measured using immunocytochemistry of thousands of mouse fibroblasts exposed to various TNF levels [15]. Further details of the method used are discussed in [16].

## RESULTS

Based on single-cell data analysis of the TNF–NF-κB signaling pathway, it has been revealed that a cell can differentiate between low and high TNF level concentrations at the input [5]. The presence of molecular noise may impede the cell’s ability to accurately discriminate the strength of the input signals and external stimulations. Consequently, it affects the cellular decision-making results [5]. As previously noted, due to numerous diseases and pathological conditions that may arise from unanticipated and altered cellular decisions, there is a keen interest in devising suitable models for the characterization and quantification of molecular signal detection factors and cellular decision-making.

To analyze the proposed cellular decision-making framework for the signaling pathway in Fig. 1, using nuclear NF-κB responses measured at the output upon receiving various TNF concentration levels at the system input, we focus on two types of decision probabilities as the desired metrics: probability of false alarm *P*_FA_, and probability of detection *P*_D_ (defined in METHODS). We evaluate the proposed model by choosing 0.0021, 0.0052, and 0.013 ng/mL as the low TNF levels (null hypothesis H_0_) vs. 0.2, 0.51, 1.3, and 50 ng/mL for high TNF levels (alternative hypothesis H_1_). For instance, Fig. 2A shows the histograms of NF-κB responses of hundreds of cells after 30 minutes of exposure to a low TNF level of 0.0052 ng/mL vs. a high TNF level of 0.2 ng/mL. Fig. 2B shows the associated univariate PDFs of the responses in Fig. 2A, including the optimal decision threshold value when *P*_FA_ = 0.1 obtained by the Neyman-Pearson theorem (details in METHODS). The vertical blue line as the optimal decision threshold is the best value that maximizes the probability of detection *P*_D_, at *P*_FA_ = 0.1. Based on the definition in the Equation (4), the right-side area of the optimal decision threshold line, shaded in light red, represents the probability of false alarm *P*_FA_ region. Also, the probability of detection *P*_D_ region is on the right side of the obtained decision threshold under the associated PDF curve of high TNF level (hypothesis H_1_), which is not shaded due to possible overlap with the *P*_FA_ region. Fig. 2C also illustrates the same PDFs for both low and high TNF levels in Fig. 2B, but includes the shaded false alarm region when *P*_FA_ = 0.7. The expansion of the shaded *P*_FA_ region is expected. This increase in threshold enlarges the *P*_D_ region as well, which means a higher *P*_D_ value.

**Figure 2.**
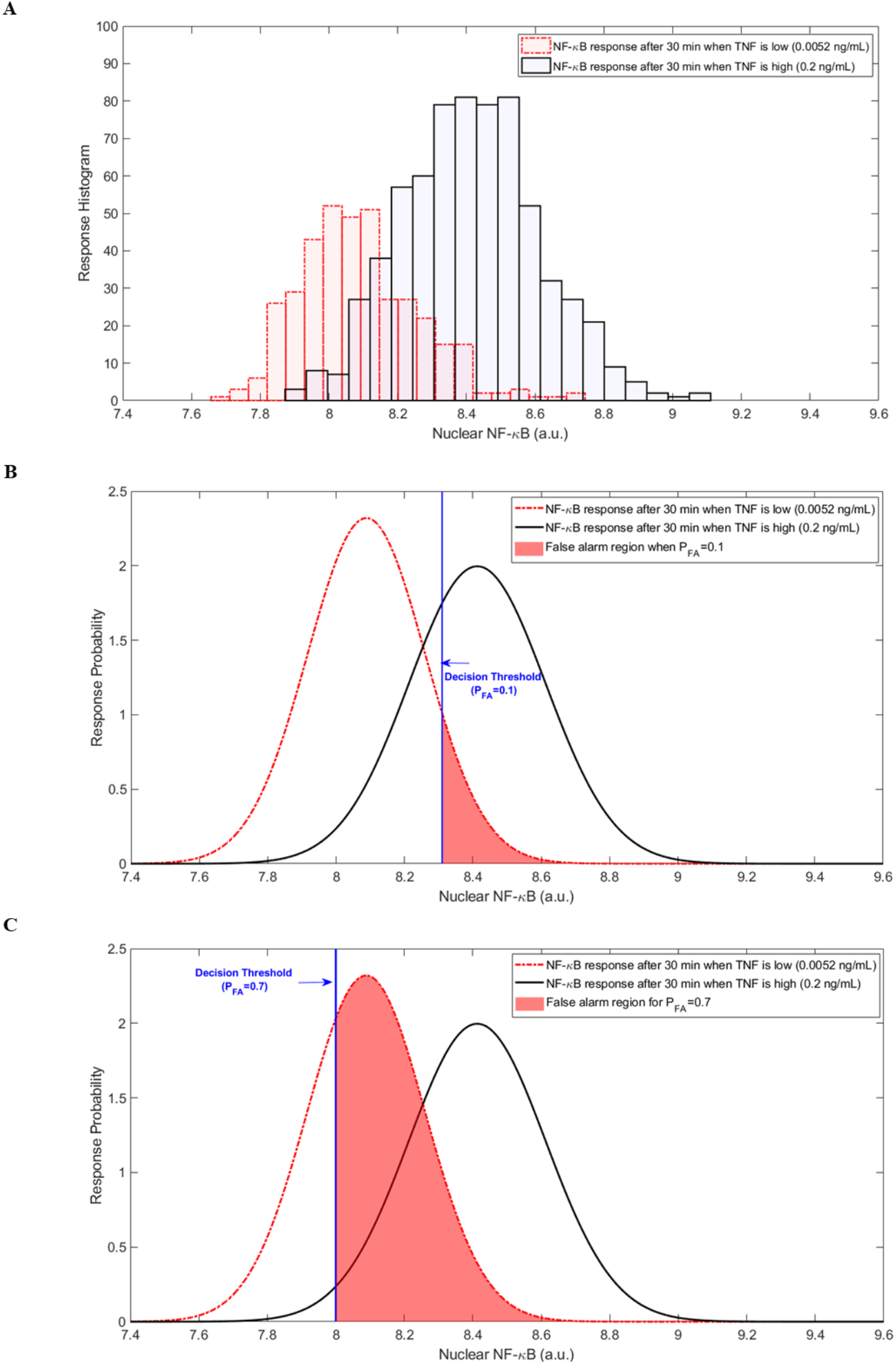
A) Histograms of measured NF-κB concentrations for low TNF 0.0052 ng/mL vs. high TNF of 0.2 ng/mL. B) Gaussian PDF of NF-κB responses to low and high TNF levels after 30 minutes of exposure including the optimal decision threshold when *P*_FA_=0.1 shown as blue vertical line and the *P*_FA_ associated region. C) Same PDFs when *P*_FA_=0.7 and its corresponding region.

As a concise demonstration of the utilized experimental data, Fig. 3A demonstrates the mean values of cells’ responses to different TNF levels in NF-κB after 30 minutes of TNF exposure. That is one of the important variables that can help to justify ROCs behaviors in Fig. 4, as well. Also, this figure tells us that as the TNF level increases in each case, the corresponding mean increases as well due to the shifting of histograms to the right in the coordinate system. Fig. 3B shows the standard deviation of cells’ responses to different TNF levels in NF-κB after 30 minutes of TNF exposure, where there are no high differences and fluctuations between various TNF levels’ standard deviations. When it comes to analyzing data, we invoke ROC (explained in METHODS) curves, here to see how the signaling system performs. Fig. 4 shows the hypothesis testing problems of 0.0052 ng/mL as the low TNF level vs. 0.2 ng/mL and 50 ng/mL as the two high TNF levels over two different time points after TNF exposure, 30 minutes and 4 hours (individually and jointly analyzed) in both wild-type and A20^−/−^ cells. First, by comparing Fig. 4C with Fig. 4A associated with wild-type cells for different high TNF levels, 50 ng/mL and 0.2 ng/mL, respectively, this suggests that there is a visible reduction in all ROCs when the high TNF level decreases from 50 ng/mL to 0.2 ng/mL, which is consistent with our earlier analysis. A lower concentration of high TNF, like 0.2 ng/mL, has a closer histogram to the low TNF of 0.0052 ng/mL than the highest high TNF, which is 50 ng/mL.

**Fig. 3.**
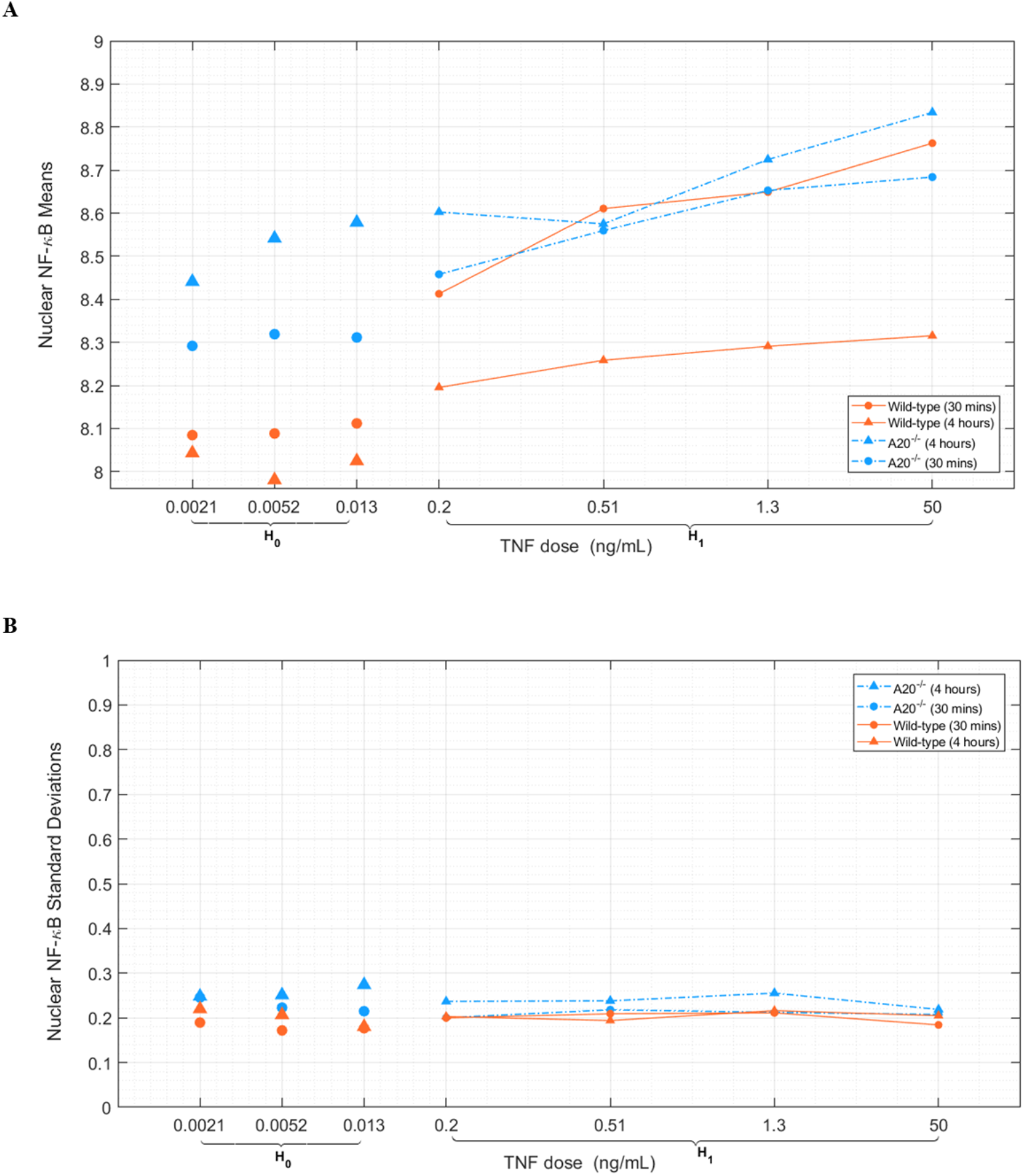
Dynamic range of responses to various TNF levels. A) Nuclear NF-κB means for various TNF levels. B) Standard deviations of Nuclear NF-κB in response to different TNF levels.

**Fig. 4.**
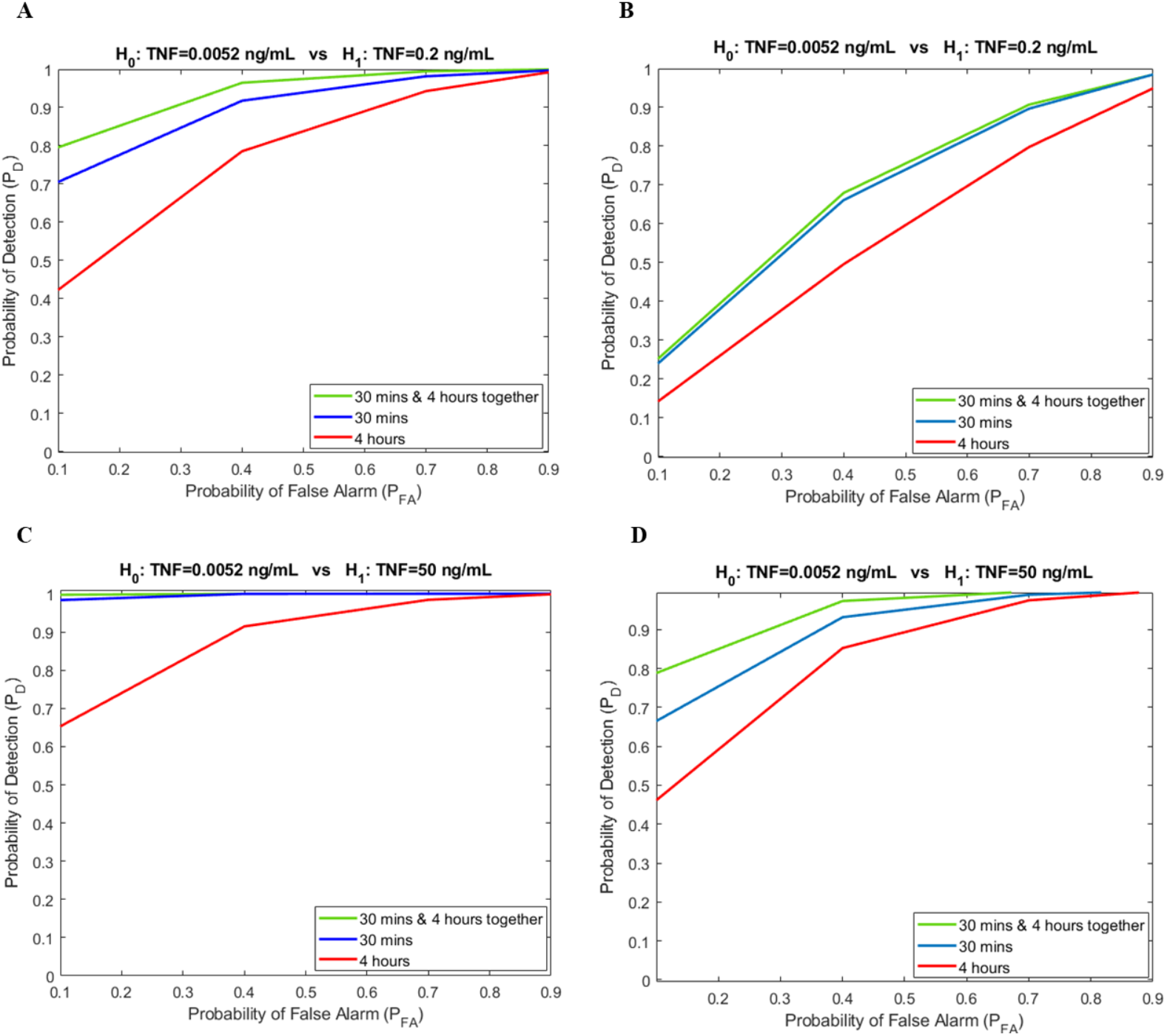
Receiver operating characteristic (ROC) curves for TNF–NF-κB signaling pathway evaluated using Neyman-Pearson theorem concepts when low TNF, H_0_ is 0.0052 ng/mL. A) Wild-type cells result for high TNF, H_1_ equal to 0.2 ng/mL. B) A20^−/−^ cells results for high TNF, H_1_ equal to 0.2 ng/mL. C) Wild-type cells result for high TNF, H_1_ equal to 50 ng/mL. D) A20^−/−^ cells results for high TNF, H_1_ equal to 50 ng/mL.

Another important observation could be about the comparison of 4-hour ROCs with 30-minute ROCs for both high TNF levels, 0.2 ng/mL and 50 ng/mL, where we can say the 4-hour curves are below the 30-minute curves due to the activation of the negative feedback over time that reduces NF-κB level after 4 hours. In A20^−/−^ cells, we expect to see the opposite behavior from wild−type. Because in the A20^−/−^ case, the negative feedback has become ineffective, and the pathway is almost linear. So, we can say *P*_D_ gets increased over time, or the *P*_D_ in 4 hours is higher than *P*_D_ in the 30-minute case. The transition from wild-type to A20^−/−^ is like shifting the corresponding histograms to the right-hand side of the coordinate system. The means figure, Fig. 3A, justifies this claim very well through a comparison of associated mean values of low and high TNF levels in the corresponding hypothesis testing. Both H_0_ and H_1_ histograms in A20^−/−^ have shifted to the right, and the means are good representative of their histograms or their PDFs. In those figures, we can clearly see that the difference between the H_0_ means and H_1_ means plays an important role in the *P*_D_ of the Neyman-Pearson decision-making framework. Because, e.g., as the means become farther away, it seems like the histograms or PDFs are well-separated, and we can say the probability of detection is higher because the PDFs or histograms are nearly well separated. Also, these can be pointed out as the opposite, as well. Those cases with lower *P*_D,_ in each case, have a closer means of H_0_ and H_1_. As we can see, analyzing both time points, 30 minutes and 4 hours together, is like a bivariate PDF that can improve the decision-making performance. The bivariate case almost always stays either very close to the one with higher *P*_D_ or at the top of each individual 30-minute and 4-hour time points, reasonably. Moreover, by comparing the above ROCs from left (Fig. 4A and 4C) to the right (Fig. 4B and 4D), cell type transition from wild-type to A20^−/−^, we always notice a reduction in all cases, 30 minutes, 4 hours, and the bivariate scenario, as well.

To better sense the excellence of bivariate analysis against univariate analysis in our cellular decision-making framework, Fig. 5 is illustrated in which for a fixed *P*_FA_ = 0.1, where low TNF is 0.0052 ng/mL, the probability of detection *P*_D_ for several high TNF levels is graphed, e.g., for wild-type cells, the bivariate *P*_D_(30 minutes & 4 hours) = 0.8, while the individual *P*_D_(30 minutes) = 0.71 and *P*_D_(4 hours) = 0.42 (Table 1). Another noteworthy impression we mentioned above is more obvious here when we compare the wild-type and A20^−/−^ cases for each high TNF levels, e.g., for the high TNF level of 0.2 ng/mL, the *P*_D_ drop is noticeable for individual time point comparison, *P*_D_(wild-type at 30 minutes) = 0.71 while *P*_D_(A20^−/−^ at 30 minutes) = 0.24. When it comes to 4-hour analysis, we can notice the same behavior, *P*_D_(wild-type at 4 hours) = 0.42 while *P*_D_(A20^−/−^ at 4 hours) = 0.14 (Table 1). Besides that, Table 2 shows the same behavior when the high TNF level is 50 ng/mL vs. the same low TNF of 0.0052 ng/mL.

**Table 1.** Decision Probabilities of wild-type and A20^−/−^ cells for TNF=0.0052 ng/mL vs. TNF=0.2 ng/mL.

| Decision Probability | Cell type | 30 minutes | 4 hours | (30 minutes & 4 hours) |
| --- | --- | --- | --- | --- |
| $P_D$<br>(at $P_{FA}=0.1$) | WT | 0.71 | 0.42 | 0.8 |
|  | A20<sup>-/-</sup> | 0.24 | 0.14 | 0.26 |

**Table 2.** Decision Probabilities of wild-type and A20^−/−^ cells for TNF=0.0052 ng/mL vs. TNF=50 ng/mL.

| Decision Probability | Cell type | 30 minutes | 4 hours | (30 minutes & 4 hours) |
| --- | --- | --- | --- | --- |
| $P_D$<br>(at $P_{FA}=0.1$) | WT | 0.98 | 0.65 | 1 |
|  | A20<sup>-/-</sup> | 0.67 | 0.46 | 0.79 |

**Figure 5.**
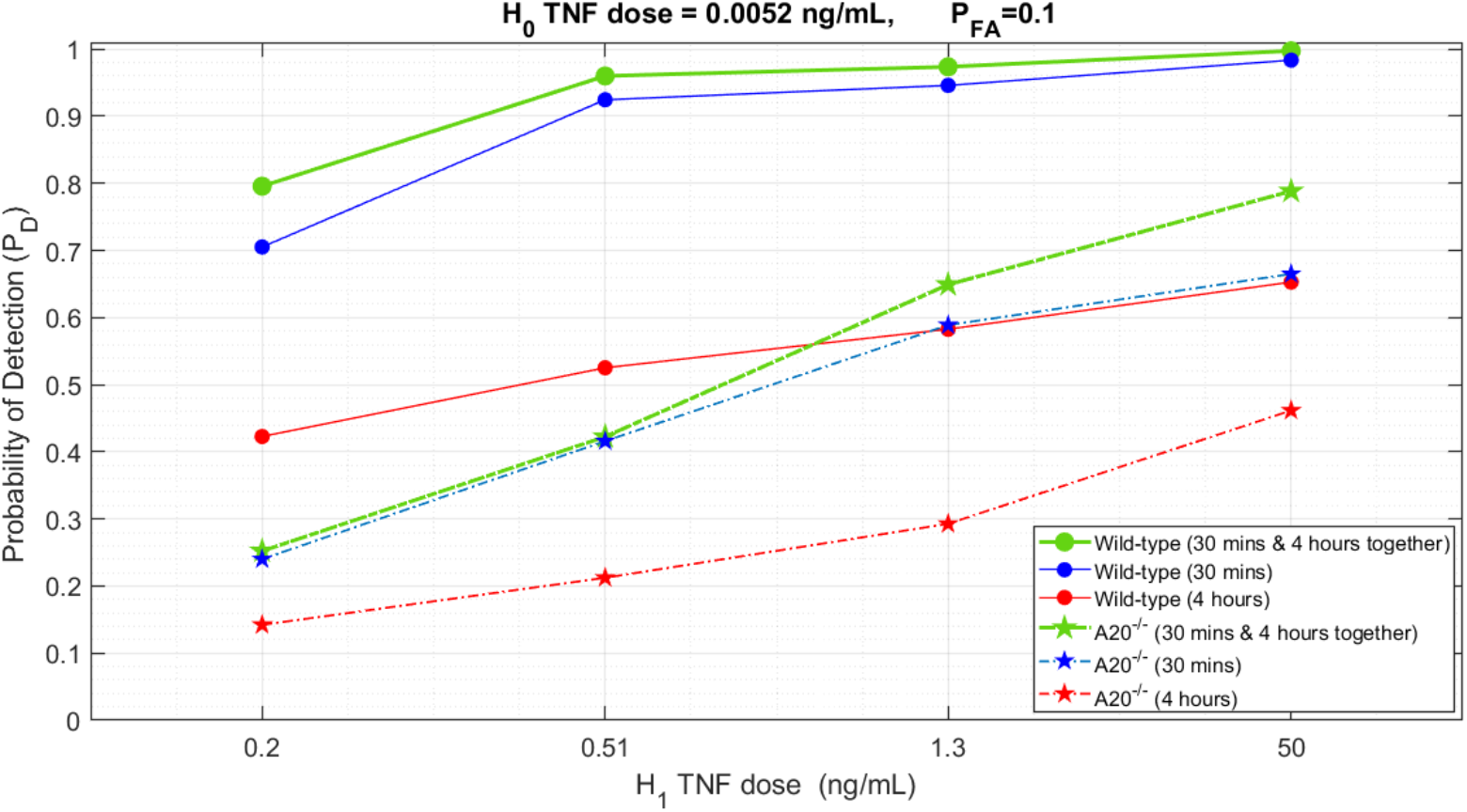
Probability of Detection vs. various TNF levels for both wild-type and A20^−/−^ cells at different time points.

These analyses are also done for another two different low TNF levels, 0.0021 ng/mL and 0.013 ng/mL, vs. the aforementioned high TNF levels, 0.2 ng/mL and 50 ng/mL, whose results are all in good agreement.

The corresponding ROCs, along with their eigenvalue decompositions to be used in the Gil-Pelaez lemma for decision-making analysis, are fully provided in SUPPLEMENTARY MATERIALS. These findings provide additional validation of the significance of this parameter within the system model, as we observed biologically meaningful outcomes.

### Analysis of Mixed Cell Populations

To further demonstrate the biological relevance of the framework, we analyzed a scenario motivated by cancer biology where tumors often contain heterogeneous populations of cells with varying A20 expression. We modeled a mixed population where a fraction β consists of A20^−/−^ cells and (1-β) are wild-type cells, with both populations exposed to either low (H_0_: 0.0052 ng/mL) or high (H_1_: 50 ng/mL) TNF concentrations. Basically H_0_: (1-β) WT + β A20^−/−^ at TNF=0.0052 ng/mL vs. H_1_: (1-β) WT + β A20^−/−^ at TNF=50 ng/mL. This scenario reflects the reality that A20^−/−^ cancer cells within a tumor microenvironment may misinterpret low TNF signals as high and trigger antiapoptotic or proliferative signaling.

Fig. 6 demonstrates the probability of detection *P*_D_ at a fixed false alarm probability *P*_FA_ = 0.1 as a function of the A20^−/−^ fraction β. As expected, detection performance decreases as the proportion of A20^−/−^ cells increases. At the boundary conditions, the results match our earlier findings: for β = 0 (pure WT), *P*_D_ values are 0.98 (30 minutes), 0.65 (4 hours), and almost 1 (bivariate); for β = 1(pure A20^−/−^), *P*_D_ decreases to 0.67, 0.46, and 0.79, respectively (Table 2).

**Figure 6.**
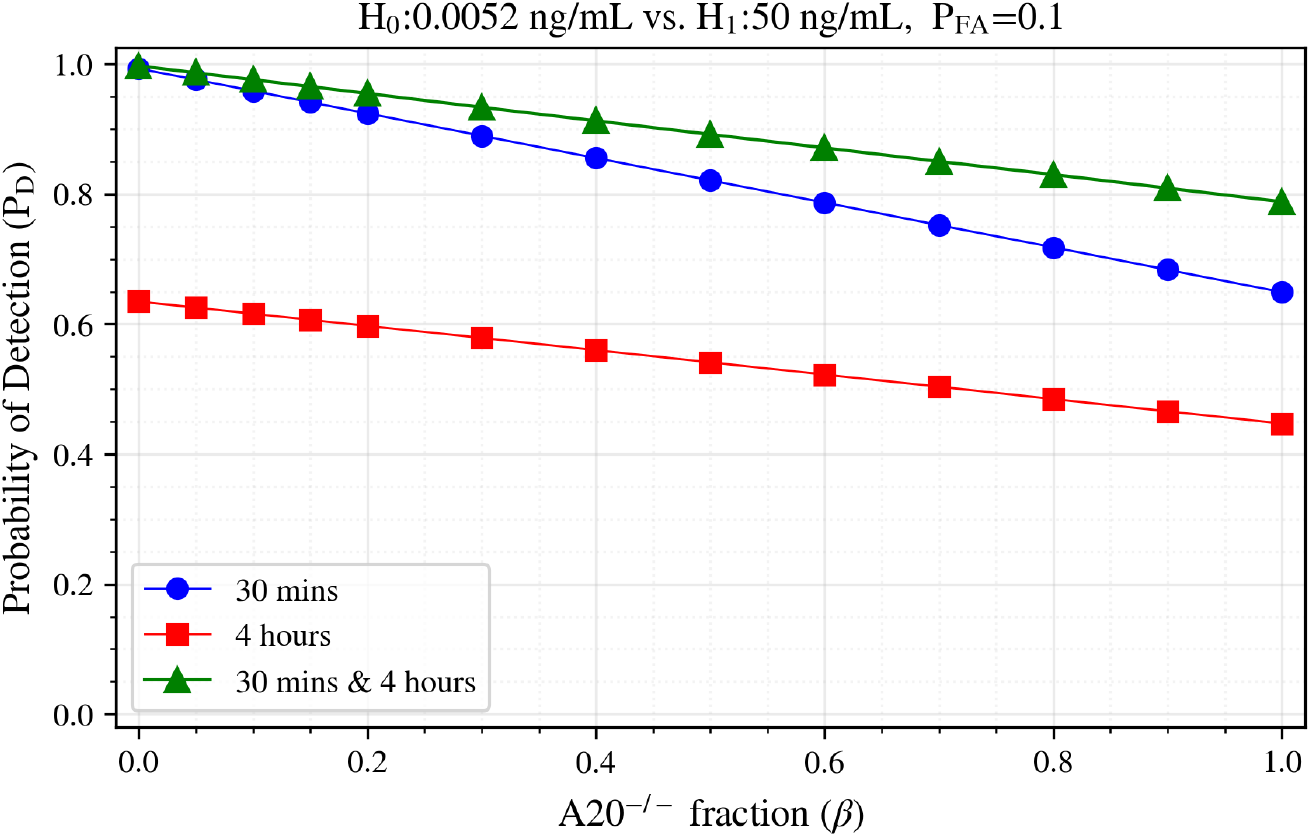
Probability of Detection *P*_D_ at *P*_FA_ = 0.1 as a function of A20^−/−^ cell fraction β for mixed populations in H_0_=0.0052 ng/mL vs. H_1_=50 ng/mL at different time points.

Fig. 7 also presents *P*_FA_ at fixed *P*_D_ = 0.8, illustrating false alarm probability at a given detection rate as the proportion of A20^−/−^ cells increases in the population. The combined 30 minutes & 4 hours analysis (green curve) maintains the false alarm rate nearly twice lower than that obtained for the best single data point (of 30 minutes). We thus observed in Figs. 6 and 7 that the bivariate analysis (green curves) consistently outperforms both univariate analyses based on single time points across the entire range of mixed compositions, demonstrating the robustness of multi-time point analysis also in heterogeneous populations.

**Figure 7.**
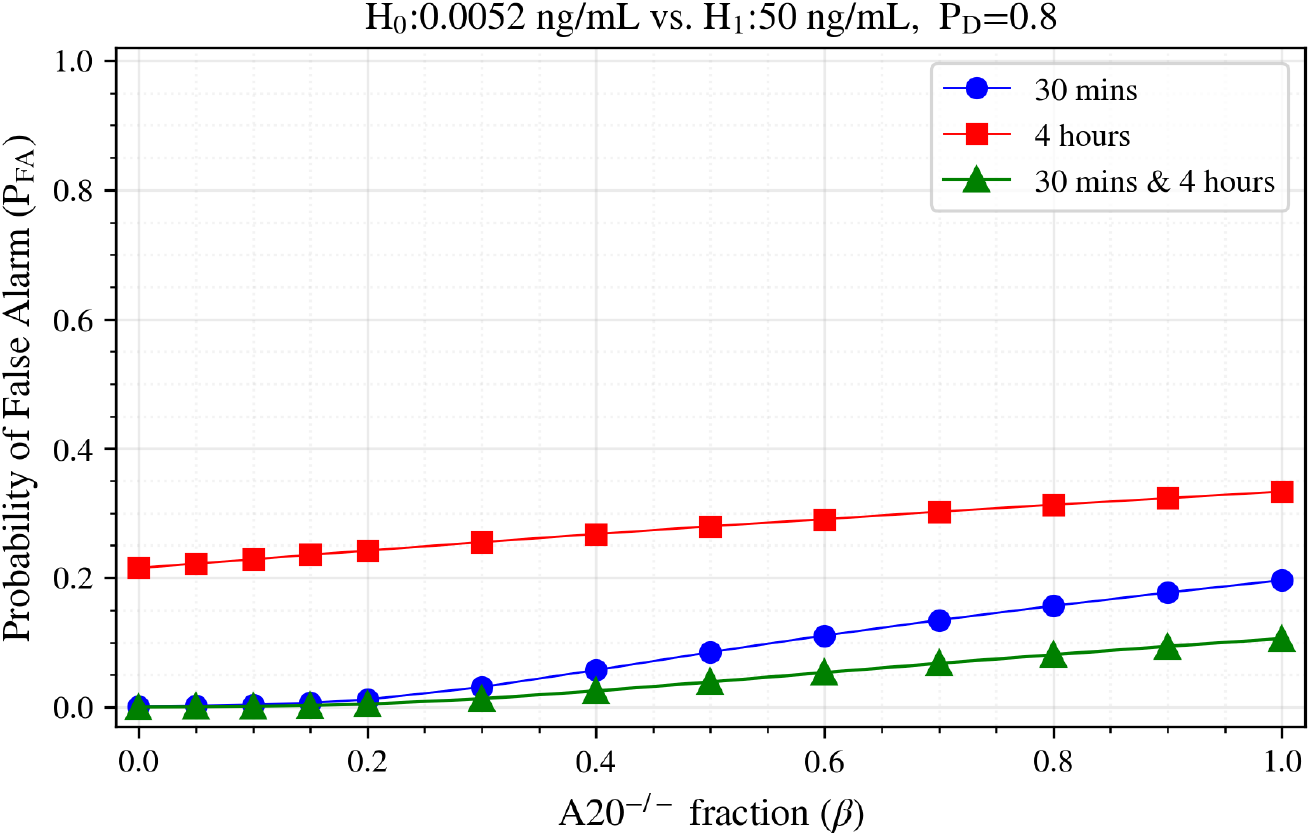
Probability of False Alarm *P*_FA_ at *P*_D_ = 0.8 as a function of A20_−/−_ cell fraction β for mixed populations in H_0_=0.0052 ng/mL vs. H_1_=50 ng/mL at different time points.

## METHODS

To enhance comprehension of the proposed model in molecular biology and cellular decision-making processes, we applied statistical signal processing and decision theory approaches to analyze the TNF–NF-κB signaling system. A cell is able to discriminate whether the TNF concentration is low or high when exposed to varying levels of TNF as an input signal [5]. However, due to potential signaling abnormalities or noise in signal transduction, the cell may respond differently to the same input, leading to incorrect decision-making. To address this challenge, we pose a binary hypothesis testing problem in which the cell should determine the true hypothesis based on the TNF input level [7].

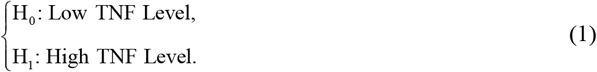

In the previous works by Habibi et al. [7], Ozen et al. [8], and Emadi et al. [9] the focus was on minimizing the probabilities of errors that a cell can make, *P*_FA_ and *P*_M_ as the statistical metrics of interest, where the first one corresponds to the probability of when a cell incorrectly determines that the input level of TNF is high when it is actually low, and the second one refers to the probability of when a cell overlooks a high level of TNF at the input. The optimal decision-making based on the maximum likelihood approach needed a knowledge of the prior probabilities of the hypotheses H_0_ and H_1_, *P*(H_0_) and *P*(H_1_), as a prerequisite to determine the optimal decision threshold and compute the error probabilities. But typically, in cell signaling studies, such information is unavailable; consequently, they were fairly assumed equal, *P*(H_0_) = *P*(H_1_) = 0.5 in previous works [7]-[9]. But in this study, we introduce a new approach, based on the Neyman-Pearson theorem, as an alternative approach for cellular decision-making studies which does not ever rely on prior knowledge of the probabilities of the hypotheses H_0_ and H_1_, *P*(H_0_) and *P*(H_1_), and aims to maximize the probability of detection *P*_D_, meaning the probability of identifying the presence of a signal when it does in fact exist [17].

Let *P*_FA_, probability of false alarm, and *P*_D_, probability of detection, be the metric decision probabilities of interest for a cell. *P*_FA_ is indeed the probability of deciding the high level of TNF at the input of the system while it is low (i.e., declaring H_1_ while H_0_ is true), and *P*_D_ is the probability of deciding the high level of TNF when in fact, it is high (i.e., declaring H_1_ while H_1_ is true), as shown below:

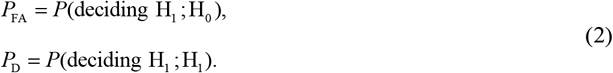

The optimal decision-making framework strives to maximize *P*_D_ for any given *P*_FA_. To better understand how it works, let vector **y**, the *N*-element observation data (as discussed later, **y** is composed of NF-κB responses), be used to form the likelihood ratio *L*(**y**) in Equation (3). It declares hypothesis H_1_when the likelihood ratio *L*(**y**) is greater than the threshold γ, and declares H_0_ otherwise, according to the Neyman-Pearson theorem [17]-[19].

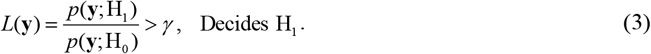

To further analyze the performance of the proposed framework, we need to compute the optimal decision threshold. First step is to define the metric parameters of interest, probability of false alarm *P*_FA_, and probability of detection *P*_D_ which are computed as follows:

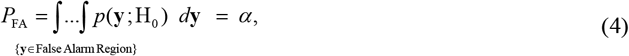

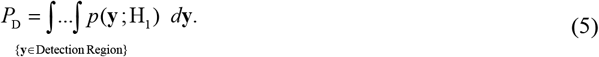

Where *p*(**y**; H_0_) and *p*(**y**; H_1_) are the conditional probability density functions (PDFs) of **y** under hypotheses H_0_ and H_1_, respectively. There is a specific γ value to be calculated for any given *P*_FA_ equal to α in Equation (4) by which the maximized *P*_D_ can be obtained as shown in Equation (5). Upon obtaining these decision probabilities, we can also take advantage of a well-known graphical representation, the Receiver Operating Characteristic (ROC), to visualize the performance of the proposed decision-making framework (Fig. 4).

The utilized experimental data are plausibly modeled by the lognormal PDFs. In other words, the natural logarithm of data, ln(data), follows the Gaussian PDF. The multivariate Gaussian PDF of the measured response quantity of data, for **y** under the hypothesis H_*i*_ is shown below [17]:

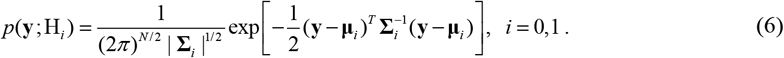

Where **y** represents the *N* ×1 column vector of the ln(data), **µ**_*i*_ is the *N* ×1 mean vector of **y** under hypothesis H_*i*_ and Σ _*i*_ is the *N* × *N* covariance matrix of y under hypothesis H_*i*_. Σ_*i*_ and 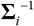 demonstrate the determinant and inverse of the matrix Σ_*i*_, respectively. And ^*T*^ denotes the transpose operator.

As mentioned earlier, to assess the performance of the proposed framework, the optimal decision threshold and the probabilities of interest in Equations (4) and (5) need to be computed in the following subsections, in a simpler case of univariate, and the general multivariate case.

### A. Univariate Decision Analysis

We use nuclear NF-κB responses of thousands of cells exposed to various TNF concentrations measured at 30 minutes and 4 hours after exposure [5]. For univariate analysis, we let y = ln (nuclear NF-κB) for each time point, individually. As mentioned earlier, a Gaussian PDF can well represent the data, so for the case of univariate analysis, when *N* = 1 in Equation (6):

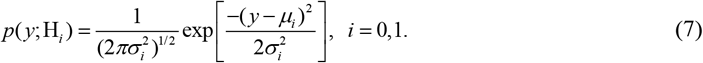

Where *μ*_*i*_ is the mean and 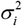 is the variance of data under *i*-th hypothesis H_*i*_. Under H_0_, TNF concentration levels 0.0021, 0.0052, and 0.013 ng/mL, and under H_1_, TNF concentration levels of 0.2, 0.51, 1.3, and 50 ng/mL were used.

To analyze the univariate decision, first, we need to define the random variable *Y* as below:

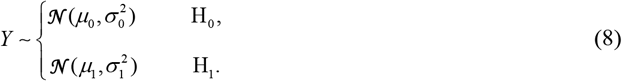

Where 𝒩 stands for the Normal (Gaussian) PDF. Equation (8) indicates that the random variable *Y* under hypothesis H_0_ has a normal distribution with mean *μ*_0_ and variance 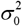. It also has a normal distribution with mean *μ*_1_ and variance 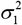, under hypothesis H_1_. Then, the transformation *X* = (*Y* − *μ*) / σ is applied to make the normal distribution 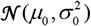 under H_0_, a normal distribution of zero mean and unit variance.

This transformation helps us in further simplification of equations and makes analytics more straightforward when it comes to the case of multivariate decision analysis. So, the hypothesis testing problem in Equation (8) for random variable *Y* turns into the new hypothesis testing problem of Equation (9) for *X*, which is actually the new transformed univariate random variable *Y*.

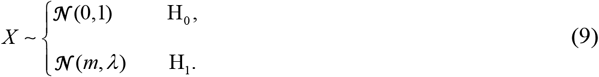

Where 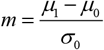 and 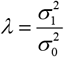.

To find the test statistic *Z* (a statistic, as a function of the data), we need to form the likelihood ratio first. As mentioned above, the importance of the transformation above will become more apparent in the next subsection **B**, when we look at *N* > 1, where finding the PDF of the test statistic is rather complicated.

The optimal decision rule declares H_1_, if:

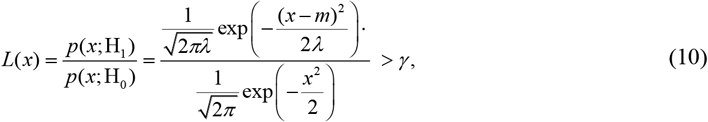

to find the optimal decision threshold for *x*, and after further algebraic simplification of the Equation (10), we obtain the following quadratic equation:

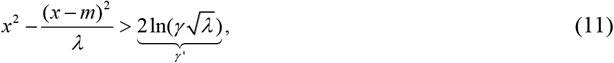

so, using the Equation (11), the test statistic can be defined as below:

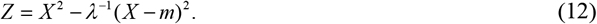

It is worth mentioning that for Gaussian class-conditionals with unequal variances 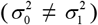, the log-likelihood ratio is quadratic in the observation, and consequently the Neyman-Pearson critical region can comprise two disjoint thresholds rather than a single boundary. We compute both algebraic roots of the quadratic equation and determine the rejection region that attains the target *P*_FA_. In practice, for most conditions in our dataset, one threshold lies in a negligible tail of the distribution where effectively no probability mass resides, reducing the decision rule to a single effective threshold. Also, for the bivariate case (*N* = 2), the *L*(**x**) = *γ* contour forms a closed boundary in the two-dimensional log-response plane (see our previous study in [9]).

To obtain the optimal decision threshold *Z*_th_ from *P*_FA_ in Equation (4), we use the Cumulative Distribution Function (CDF) of the test statistic *Z, F*_*Z*_ (*z*):

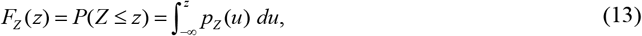

So, the decision probabilities of interest in the Equations (4) and (5) can be calculated using Equation (13), as follows:

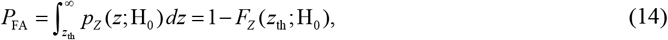

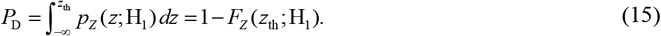

Where the *F*_Z_ (*z*_*th*_; H_*i*_) is the conditional CDF of the test statistic *Z* given hypothesis H_*i*_. As discussed earlier, for any given *P*_FA_ = α, the *Z*_th_ can be obtained from the Equation (14), but first the according *F*_Z_ (*z*) needs to be calculated.

To calculate *F*_Z_ (*z*) through the Gil-Pelaez lemma [20][21], the characteristic function of the test statistic *Z* is required:

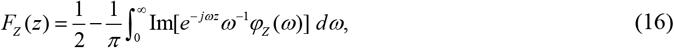

where the Im[*x*] is the imaginary part of *x* and 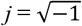.

As mentioned above, the characteristic functions of the test statistic *Z* are needed for computing *P*_FA_ and *P*_D_. Then, *φ*_*Z*_ (*ω*) under hypothesis H_0_ and H_1_ can be obtained as below:

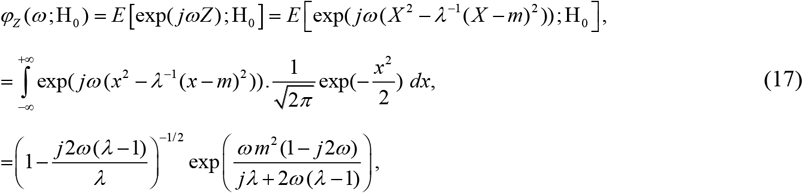

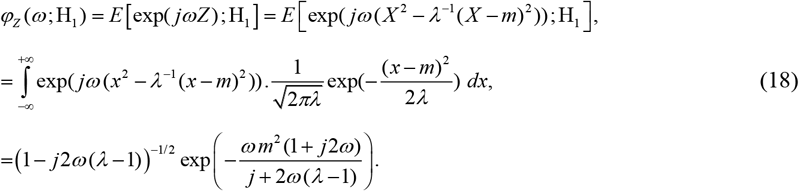

Here, *E*[*x*] denotes the expected value of *x*. This route is especially useful when *N* > 1. In principle, one could obtain *F*_Z_(*z*) by integrating the PDF *p*_*z*_(*z*), but for our test statistic *Z*, the PDF has no simple closed form when *N* > 1 (see Subsection B). Instead, we compute *F*_Z_(*z*) numerically by using the characteristic function from Equations (17) and (18) together with the Gil-Pelaez inversion in Equation (16). With *F*_Z_(*z*) in hand, *P*_FA_ and *P*_D_ can be computed using Equations (14) and (15).

### B. Multivariate Decision Analysis

In the general case, we further expand the study to multivariate decision analysis when *N* > 1, e.g., when *N* = 2, the decision probabilities of interest when NF-κB responses are analyzed for the concurrent case of two different time points of 30 minutes and 4 hours after TNF exposure in the RESULTS section. To compute the *P*_FA_ and *P*_D_ for this case, we need to take all the steps of univariate decision analysis (subsection **A**). First, for the *N*-element vector **Y** below, we use the multivariate Gaussian PDF of Equation (6) to form the likelihood ratio for the hypothesis testing problem in Equation (19).

So, for the multivariate random variable *Y*, we have:

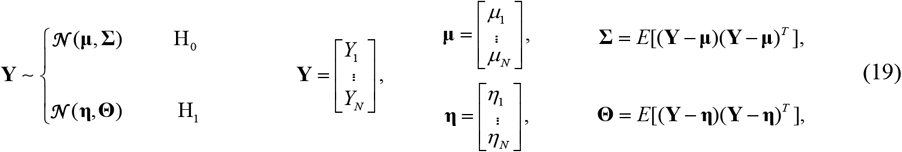

where **µ** is the *N* ×1 mean vector and **Z** is the *N* × *N* covariance matrix under hypothesis H_0_. Also, **η** is the *N* ×1 mean vector and **Θ** represents the *N* × *N* covariance matrix 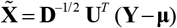 under hypothesis H_1_.

As discussed earlier, to normalize the PDF under H_0_, making it a zero-mean and unit-variance Gaussian PDF, we do a transformation that obviously affects the PDF under H_1_, as well. First, we start with an eigenvalue decomposition of Σ, the covariance matrix under hypothesis H_0_ as follows [17]:

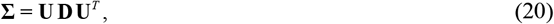

where **U** is the orthonormal matrix of eigenvectors, and **D** is the diagonal matrix of eigenvalues of the covariance matrix Σ. To transform, the variable **Y** is renormalized using two main steps; first, the mean of the PDF under hypothesis H_0_ is subtracted to obtain (**Y** − **µ**). Then de-correlating the PDF under hypothesis H_0_, **U**^*T*^ (**Y** − **µ**). And then the PDF under hypothesis H_0_is whitened, **D**^−1/2^ **U**^*T*^ (**Y** − **µ**), where **D**^−1/2^ is the diagonal matrix of the square root of the reciprocal of the eigenvalues along its diagonals. Clearly, upon doing all the processes mentioned above, the mean and covariance matrix of the PDF under hypothesis H_1_ will be transformed accordingly, as below:

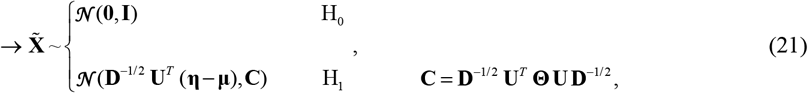

where **C** is the new covariance matrix of the PDF under hypothesis H_1_ in the Equation (21), which is determined by the two original covariance matrices. The next step would be applying another transformation to reach the new variable **X**, as the desired transformed version of variable **Y**. The new covariance matrix **C** of the transformed 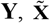 under hypothesis H_1_ can be decomposed into its eigenvectors and eigenvalue matrices as **C** = **V Λ V**^*T*^ where **V** is the orthonormal matrix of eigenvectors, and **Λ** is the diagonal matrix of eigenvalues of the covariance matrix **C**. Then, **X** is made by de-correlating the PDF under hypothesis H_1_ in Equation (21), 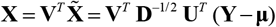. Therefore, **X** as shown below, has a zero-mean and unit-variance Gaussian PDF under H_0_, and under H_1_, a new Gaussian PDF whose mean and variance are **m** and **Λ**, respectively.

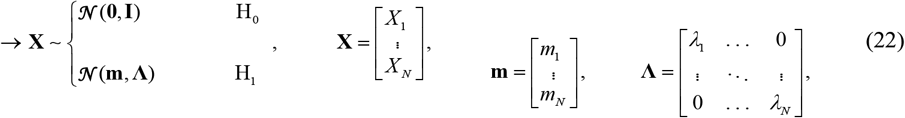

where **m** = **V**^*T*^ **D**^−1/2^ **U**^*T*^ (**η** − **µ**) is the new mean, and Λ is the new covariance matrix under hypothesis H_1_ in Equation (22).

As mentioned earlier, such a transformation can help us in further analytical simplifications. The optimal likelihood-based decision framework for variable **Y** in Equation (19) decides H_1_, if:

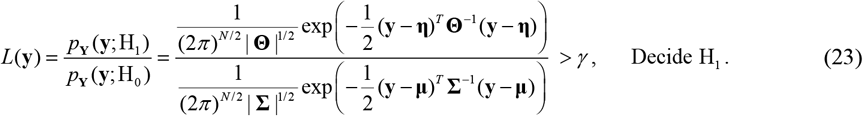

But the optimal likelihood-based decision rule for **X**, the transformed variable **Y**, declares H_1_, if:

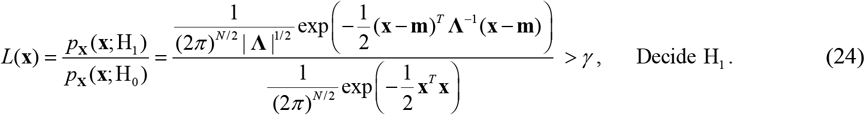

Similar to the case of univariate decision analysis (subsection **A**), after some simplifications, the test statistic *Z* can be obtained as follows:

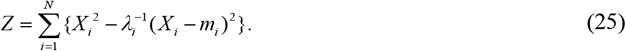

and the binary hypothesis testing problem for the multivariate decision-making framework can be defined below:

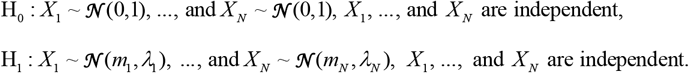

The next step is finding the PDF of the test statistic *Z*, then computing the decision probabilities *P*_FA_ and *P*_D_. To do so, the characteristic function of the data under each hypothesis is needed. For the data under hypothesis H_0_, we have:

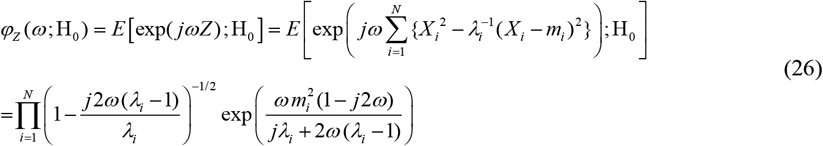

and the characteristic function of the data hypothesis H_1_, will be calculated as below:

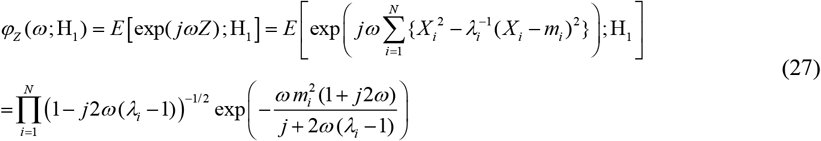

According to Equations (3)-(5), and similar to univariate decision analysis, *F*_*Z*_(*z*) can be numerically obtained using the characteristic function of *Z* in Equations (26) and (27), together with the Gil-Pelaez lemma. Eventually, we can find *P*_FA_ and *P*_D_.

Here, in the case of *N* = 2, we consider two time points, 30 minutes and 4 hours, to be simultaneously considered as the two desired variables for analyzing the decision-making framework using the experimental data set. The eigenvalue decompositions of all possible conditions analyzed here are provided in the SUPPLEMENTARY MATERIALS section.

Also, this approach can be developed to larger cellular signaling networks, including different inputs like secondary messengers or ligands, and the output could be various transcription factors [22].

## CONCLUSION

The introduced approach in this work proposes a set of statistical signal processing and decision theoretic methods and metrics to model and measure the cellular decision-making errors in a TNF–NF-κB signaling system, under normal conditions and abnormality, which can be caused by noise or signaling uncertainties. This study showed biologically relevant and meaningful results that agreed with the functionality of the above-mentioned signaling pathway. The important transcription factor NF-κB regulates many genes, controls cell fate, and is involved in apoptosis and the transition of an antiviral state. So, it is highly crucial for the cell to make an accurate decision in the information transmission of TNF–NF-κB signaling pathway. Here, we focused on such an important cell signaling system for which we presented univariate and bivariate methods to analyze the cell decision-making process and outcomes using Neyman-Pearson theorem, a statistical signal processing method which has the advantage of no need to know prior probabilities, whereas previous works by Habibi et al., Ozen et al., and Emadi et al. [7]-[9] require the knowledge of prior probabilities. The signaling pathway could transmit the information of changes in TNF concentration level from 0.0021 ng/mL to 50 ng/mL. The mean estimates in Fig. 3A are derived from (and represent) the histograms in Fig. 2A. The difference between the means of H_0_ and H_1_ plays a crucial role in the probability of detection *P*_D_ of the proposed model. When the means are further apart, the histograms appear more separated, indicating a higher probability of detection. Conversely, cases with lower *P*_D_ show closer means between H_0_ and H_1_.

A key finding is that when the TNF signal is strong in wild-type cells, the signal detection probability is higher compared to the weak TNF signal detection. The univariate results in Fig. 4 demonstrate that in wild-type cells, the system outperforms A20^−/−^ cells in both time points, 30 minutes and 4 hours. Analyzing receiver operating characteristic (ROC), as a measure of system performance, indicates that both time points, 30 minutes and 4 hours, together, which can be considered as a joint or bivariate case, enhances the performance of the decision-making framework. The bivariate case consistently remains either very close to the one with higher *P*_D_ or above both univariate scenarios, the 30 minutes and 4 hours. Furthermore, when comparing the ROC curves from left to right panels (cell transition from wild-type to A20^−/−^ in Figs. 4A-4D), a reduction in performance is observed in all cases: 30 minutes, 4 hours, and the bivariate case. This advantage persists even in heterogeneous cell populations containing mixed wild-type and A20^−/−^ cells, demonstrating the robustness of the multivariate approach in biologically realistic scenarios (Figs. 6 and 7).

These findings are also consistent with biological experiments, where A20 deficiency can potentially lead to chronic inflammation or even contribute to the development of cancer. Overall, the introduced decision-theoretic model can be advantageous for understanding the performance of cellular decision-making processes without any knowledge of the stimuli’s prior probabilities that are not biologically available.

## DATA AND CODE AVAILABILITY

The dataset generated and analyzed during this study is available from the corresponding author upon reasonable request. Also, the custom analysis code and scripts are available on the First author’s GitHub (https://github.com/aliemadi21/NP_Cellular_Decision_Making).

## ACKNOWLEDGEMENTS

TL acknowledges support from the National Science Centre, Poland (ncn.gov.pl), grant 2023/51/B/NZ6/02951.

AL is supported by the NSF grant MCB-2231765 and NIH award R01 GM123011.

## AUTHOR CONTRIBUTIONS

Conceptualization, A.A.; methodology, A.A.; software, A.E.; validation, A.E.; formal analysis, A.E.; investigation, A.E.; resources, A.L. and A.A.; data curation, A.L.; writing—original draft preparation, A.E.; writing—review and editing, A.E., T.L., A.L., and A.A.; visualization, A.E.; supervision, A.A.; project administration, A.A.

All authors have read and agreed to the published version of the manuscript.

## SUPPLEMENTARY MATERIALS

**Figure S1.**
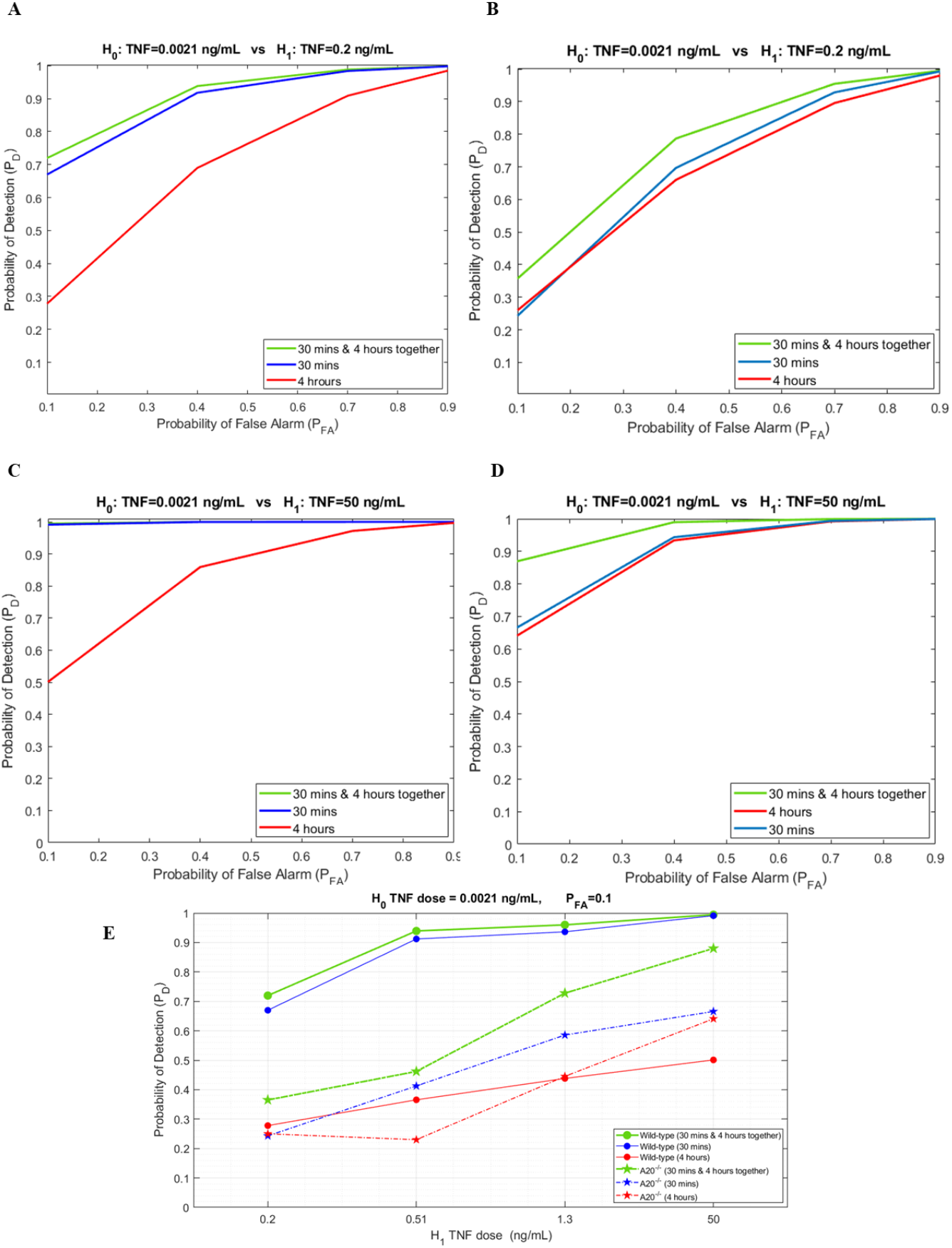
Receiver operating characteristic (ROC) curves for TNF– NF-κB signaling pathway evaluated using Neyman-Pearson theorem when low TNF H_0_ is 0.0021 ng/mL. A) Wild-type cells result for high TNF H_1_ equal to 0.2 ng/mL. B) A20^−/−^ cells results for high TNF H_1_ equal to 0.2 ng/mL. C) Wild-type cells result for high TNF H_1_ equal to 50 ng/mL. D) A20^−/−^ cells results for high TNF H_1_ equal to 50 ng/mL. E) Probability of Detection vs. various TNF levels for both wild-type and A20^−/−^ cells at different time points.

**Figure S2.**
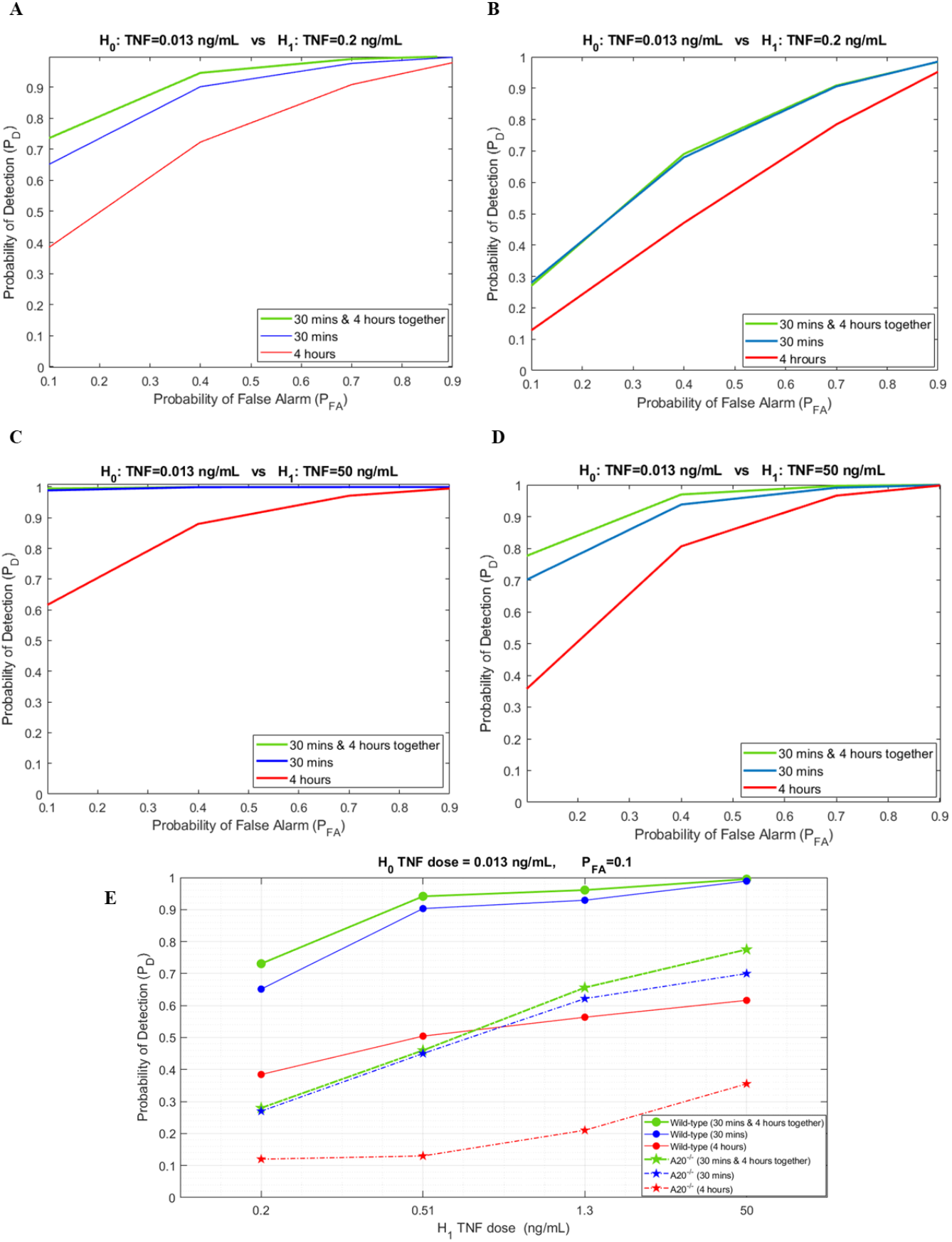
Receiver operating characteristic (ROC) curves for TNF– NF-κ B signaling pathway evaluated using Neyman-Pearson theorem when low TNF H_0_ is 0.013 ng/mL. A) Wild-type cells result for high TNF H_1_ equal to 0.2 ng/mL. B) A20^−/−^ cells results for high TNF H_1_ equal to 0.2 ng/mL. C) Wild-type cells result for high TNF H_1_ equal to 50 ng/mL. D) A20^−/−^ cells results for high TNF H_1_ equal to 50 ng/mL. E) Probability of Detection vs. various TNF levels for both wild-type and A20^−/−^ cells at different time points.

**Mean Vectors and Covariance Matrices after Eigenvalue Decomposition for Various Scenarios**

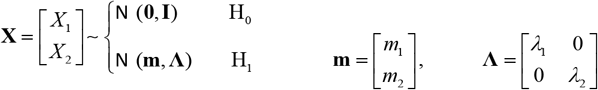

**A) Wild-type**

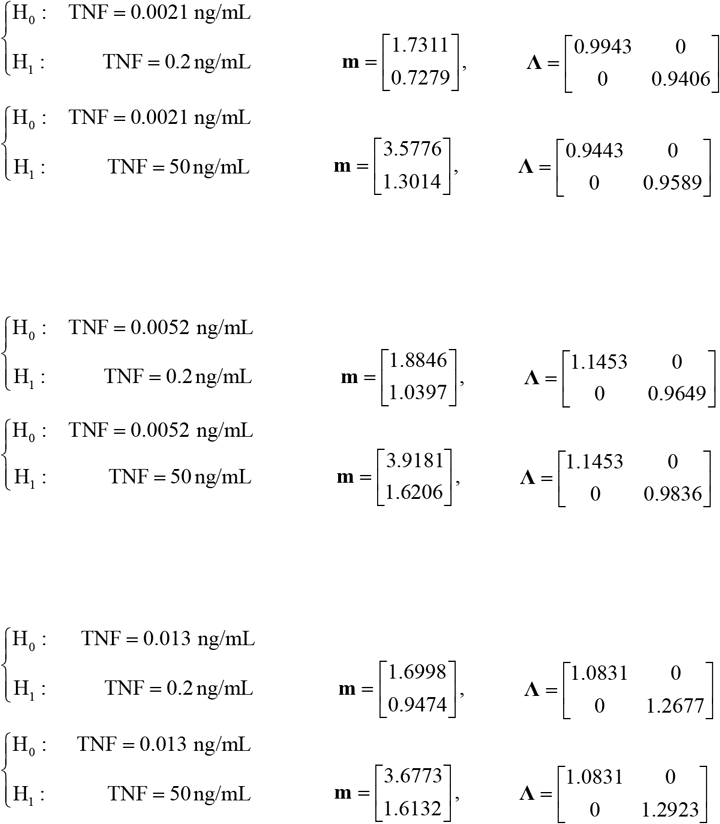

**B) A20**^**−/−**^

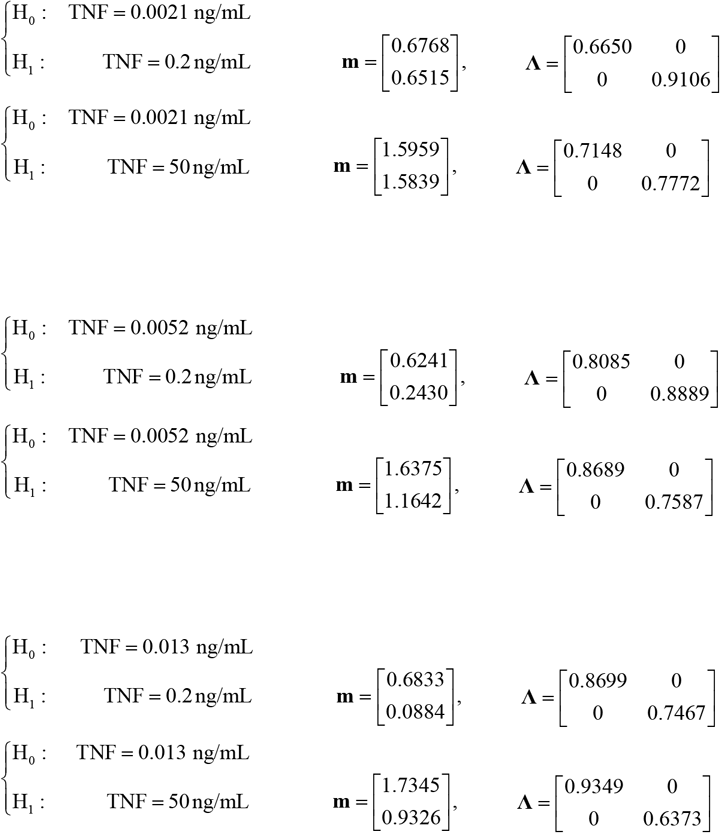

## REFERENCES

1. A. Emadi, M. Ozen, A. Abdi, A hybrid model to study how late long-term potentiation is affected by faulty molecules in an intraneuronal signaling network regulating transcription factor CREB, Integr Biol 14 (5) (2022) 111–125.

2. A. Emadi, A. Abdi, A study of how abnormalities of the CREB protein affect a neuronal system and its signals: Modeling and analysis using experimental data, in the proceedings of IEEE Signal Processing in Medicine and Biology Symposium (SPMB) (2022) 1–6.

3. A. Emadi, A. Abdi, M. Migliore, et al, A hybrid model to explore how a hippocampal CA1 neuronal network is affected by faulty molecules of an intraneuronal CREB signaling network, in the proceedings of IEEE Signal Processing in Medicine and Biology Symposium (SPMB) (2024) 1–6.

4. G. Balázsi, A. van Oudenaarden, J. J. Collins, Cellular decision making and biological noise: from microbes to mammals, Cell 144 (6) (2011) 910–925.

5. R. Cheong, A. Rhee, C. J. Wang, et al., Information transduction capacity of noisy biochemical signaling networks, Science 334 (6054) (2011) 354–358.

6. S. Tay, J. J. Hughey, T. K. Lee, et al., Single-cell NF-kappaB dynamics reveal digital activation and analogue information processing, Nature 466 (7303) (2010) 267–271.

7. I. Habibi, R. Cheong, T. Lipniacki, et al., Computation and measurement of cell decision making errors using single cell data, PLoS Comput Biol. 13 (4) (2017) 1–17.

8. M. Ozen, T. Lipniacki, A. Levchenko, et al., Modeling and measurement of signaling outcomes affecting decision making in noisy intracellular networks using machine learning methods, Integr Biol. 12 (5) (2020) 122–138.

9. A. Emadi, T. Lipniacki, A. Levchenko, et al., Single-Cell measurements and modeling and computation of decision-making errors in a molecular signaling system with two output molecules, Biology 12 (12) (2023) 1461.

10. A. Levchenko, Genetic diseases: How the noise fits in, Curr Biol. 33 (6) (2023) 228–230.

11. O. Micheau, J. Tschopp, Induction of TNF receptor I-mediated apoptosis via two sequential signaling complexes, Cell 114 (2) (2003) 181–190.

12. L. Catrysse, M. Farhang Ghahremani, L. Vereecke, et al., A20 prevents chronic liver inflammation and cancer by protecting hepatocytes from death, Cell Death Dis. 7 (6) (2016) e2250.

13. S. G. Hymowitz, I. E. Wertz, A20: from ubiquitin editing to tumour suppression, 10 (5) (2010) 332–341.

14. M. Kato, M. Sanda, I. Kato, et al., Frequent inactivation of A20 in B-cell lymphomas, Nature 459 (7247) (2009) 712–716.

15. E. G. Lee, D. L. Boone, S. Chai, et al., Failure to regulate TNF-induced NF-κB and cell death responses in A20-deficient mice, Science 289 (5488) (2000) 2350–2354.

16. R. Cheong, C. J. Wang, A. Levchenko, High content cell screening in a microfluidic device, Mol Cell Proteomics 8 (3) (2009) 433–442.

17. S. M. Kay, Fundamentals of Statistical Signal Processing: Detection Theory, PTR Prentice-Hall, New Jersey, 1998.

18. R. O. Duda, P. E. Hart, D. G. Stork, Pattern Classification, John Wiley & Sons, New York, 2001.

19. H. L. Van Trees, K. L. Bell, Z. Tian, Detection, Estimation and Modulation Theory, Part I: Detection, Estimation, and Filtering Theory, 2nd edn., Wiley, New Jersey, 2013.

20. V. Witkovský, Computing the distribution of a linear combination of inverted gamma variables, Kybernetica 37 (1) (2001) 79–90.

21. J. Gil-Pelaez, Note on the inversion theorem, Biometrika 38 (1951) 481–482.

22. A. Rhee, R. Cheong, A. Levchenko, Noise decomposition of intracellular biochemical signaling networks using nonequivalent reporters, Proc Natl Acad Sci U S A 111 (48) (2014) 17330–17335.

